# Prokaryotic microbial diversity analysis and metabolic function prediction of salt lakes on the Qinghai-Tibet Plateau

**DOI:** 10.1101/2023.08.10.552822

**Authors:** Man Zhang, Yongzhen Li, Derui Zhu, Jiangwa Xing, Qifu Long, Guoping Shen

## Abstract

The Dong Taijinar Salt Lake (DT) and Xi Taijinar Salt Lake (XT) have been widely studied as mineral-rich areas. However, little is known about the composition and distribution of the microbial communities in these two lakes. In this study, metagenomics sequencing was used to analyze the diversity and potential functions of the microbial communities in DT and XT. According to our report, the salinity of DT (332.18–358.30 g/L) was 10 times higher than that of XT (20.09–36.83 g/L). Interestingly, the dominant domain in DT was Archaea (96.16%), while that in XT was Bacteria (93.09%). The distribution of *Bacteria* in the DT revealed 33 phyla and 1717 genera. The dominant genus in DT was *Marinobacillus*, which was positively correlated with total phosphorus content. There were four main phyla and 153 genera identified in the *Archaea* of DT. The most abundant *Archaea* genera in DT were *Natronomonas* (24.61%) and *Halorubrum* (23.69%), which were mainly positively correlated with the Na^+^, Ca^2+^, and Cl^−^ contents. Similarly, there were 33 phyla and 1906 genera of *Bacteria* in XT, and *Loktanella* was the dominant genus. The archaeal taxonomy in XT mainly included four phyla and 149 genera. *Proteobacteria* and *Euryarchaeota* were the most abundant bacterial and archaeal phyla in the two salt lakes. Analysis of the halophilic mechanisms of the microorganisms identified in these two salt lakes revealed that the *Bacteria* in XT preferred to synthesize compatible solutes, whereas the *Archaea* in DT preferred a “salt-in” adaptation strategy in salt-stressed environments.

**IMPORTANCE:** The Qinghai-Tibet Plateau is the origin of many lakes and mountains in China. Among them, the Dong Taijinar and Xi Taijinar salt lakes are important biological resources with unknown microbial community compositions and functional potentials. The results of this study revealed significant differences in the distribution of *Bacteria* and *Archaea* between the two salt lakes. Salinity mainly drives lower biodiversity and restricted bacterial growth and metabolism in the high-salinity and near-saturated Dong Taijinar Salt Lake. This study not only identifies the key microorganisms in two penetrating salt lakes, but also provides insights into the mechanisms of salinity tolerance and the unknown ecological functions of microorganisms in extreme environments.

## BACKGROUND

Hypersaline salt lakes and salt marshes are formed in arid or semi-arid climates when evaporation exceeds precipitation and production is continuously concentrated (1, 2). The Qaidam Basin, located in the northeast Qinghai-Tibet Plateau, is the largest contiguous salt marsh in China (3). Dong Taijinar Salt Lake (abbreviated as DT) and Xi Taijinar Salt Lake (abbreviated as XT) are two typical penetrating salt lakes in the Qaidam Basin that share similar topographic and climatic features (Fig. 1). They both have an altitude of approximately 2628 m, and together they cover an area of approximately 100 km^2^ (4, 5). It is reported that DT is a major lithium salt producing area, which together with XT and Qarhan Salt Lake provides rich lithium resources for China and even the world (6, 7). The rich boron, potassium, and lithium resources of DT and XT are mainly due to the steady replenishment by the Nalenggler catchment (3, 8). The salinities of DT and XT, as measured by Zheng et al. in 2010, were 332.0 g/L and 338.0 g/L, respectively, which lie between the deep brines in the western and northern parts of the Qaidam Basin (4, 9, 10). Moreover, both lakes have the same brine properties, with magnesium sulfate (MgSO_4_) being the main chemical present. Interestingly, the gap in salinity between these two salt lakes has been increasing in recent years due to the salinity decreasing in XT (11, 12). The extreme decrease in salinity of XT is likely to have been caused by multiple climatic and natural factors (7).

**Figure 1.**
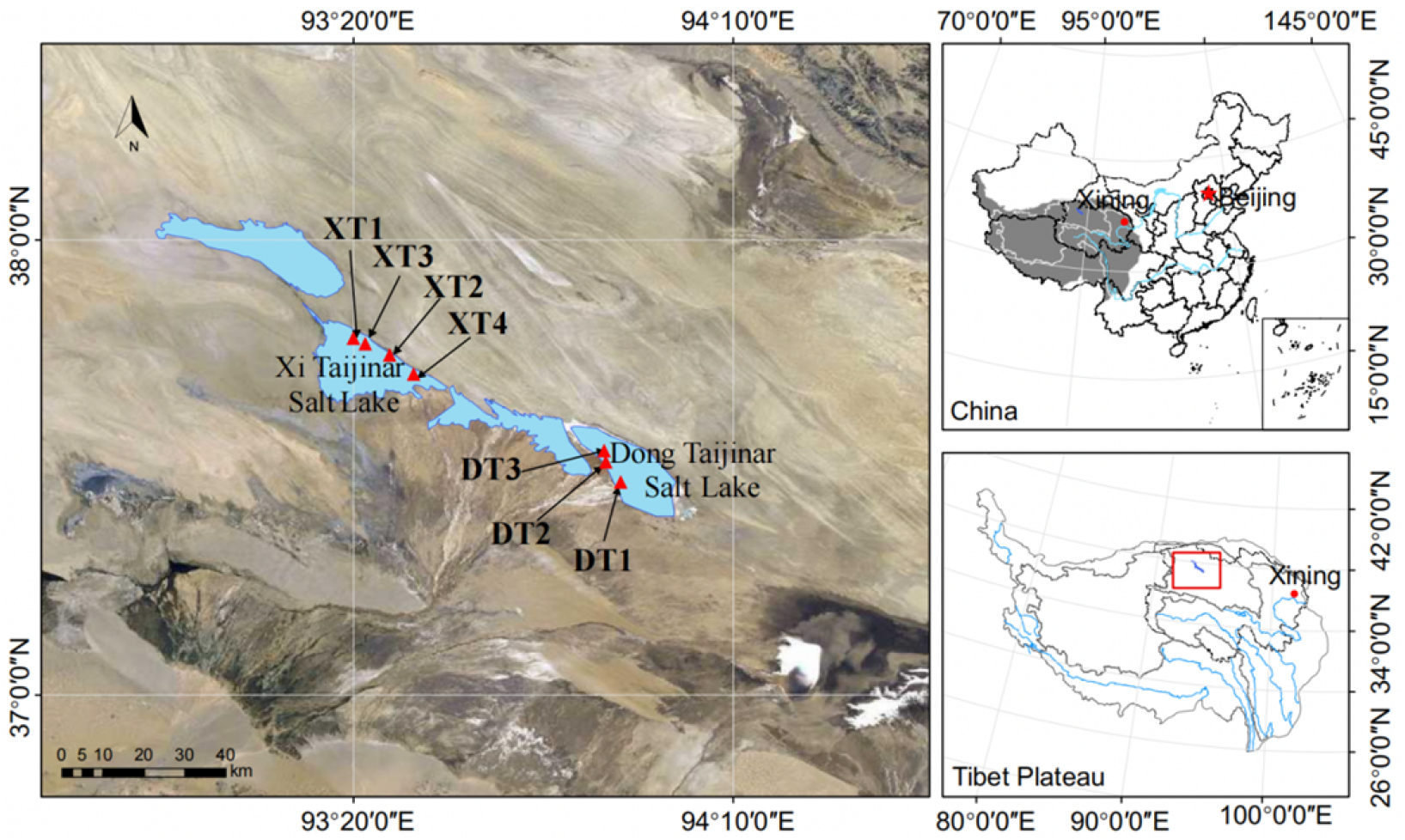
Location of the DT and XT samples. The samples from DT were obtained from the west side of the lake, and the samples from XT were obtained from the east side of the lake.

The influences of salinity on microbial diversity, community structure, and metabolic activities are significant (13). Different salinities can still be present in neighboring salt lakes, even when they are supplied from the same water source. Bishav et al. found that the difference in salinity between the northern and southern regions of the hypersaline Great Salt Lake led to species diversity (14). Plitvice lakes with rich microbial communities were attributed to rare network ecosystems of varying salinity composed of several different landforms(15). Perhaps the difference in microbial diversity due to different salinities in the same area is a unique barrier system. Therefore, extreme environments of DT and XT may also bring distinctive microbial types. To the best of our knowledge, however, few studies have reported the functional distribution and presence of microorganisms in the hypersaline lakes of DT and XT, whereas most studies have focused on the human environment and salt production. Given the unique extreme environment of salt lakes, a high saline lake provides an excellent research opportunity to explore the mechanism of salt tolerance and the function of salt-tolerant microorganisms. Metagenomic analysis techniques have been successfully applied to the structure and function of communities for applications in clinical diseases, salt lakes, reservoirs, and oceans (16–18). Compared with 16S rRNA gene analysis, metagenomic analysis not only identifies and mines most prokaryotes that are currently considered unculturable in the laboratory, but also provides a more detailed environmental gene pool and potentially relevant functions of microbial communities (19). Here, we first report the characteristics of microbial community diversity in DT and XT using metagenomics sequencing. The main aims of this study were to 1) identify key microbial players in XT and DT; 2) understand the unknown ecological functions of species in extreme environments; and 3) explore the salt-tolerance mechanisms of microorganisms in hypersaline lakes.

## RESULTS

### Environmental factors in XT and DT

The hydrochemical characteristics of DT and XT are closely related to the inflow stream (3). The intense water exchange in the NLGL in the wet season has a great effect on the brine concentration (7). Environmental parameters, including pH, total salinity (TS), and ionic composition were measured for the two salt lakes (Table 1). The TS of XT was in the range of 20.09–36.83 g/L, and the TS of DT was approximately 332.18–358.30 g/L. The total organic carbon (TOC) content in DT ranged from 8.3 mg/L to 146 mg/L, and that in XT ranged from 4.6 mg/L to 52 mg/L. Similarly, the total nitrogen (TN) in DT and XT varied. The major cations, including potassium (K^+^), sodium (Na^+^), calcium (Ca^2+^), and magnesium (Mg^2+^), and anions, including chloride (Cl^−^) and sulfate (SO_4_^2−^), differed significantly between the two salt lakes and showed enrichment in DT compared with XT. In addition, the pH of samples in DT and XT was 7.9 and 7.0, respectively.

**Table 1.** Sampling record and Environmental conditions^ac^.

| Sample name | N | E | H | T | Na<sup>+</sup> | K<sup>+</sup> | Ca<sup>2+</sup> | Mg<sup>2+</sup> | Cl<sup>-</sup> | SO<sub>4</sub><sup>2-</sup> | HCO<sub>3</sub><sup>-</sup> | TS | TN | TP | TOC | pH |
| --- | --- | --- | --- | --- | --- | --- | --- | --- | --- | --- | --- | --- | --- | --- | --- | --- |
| DT1 | 37°28'3" | 93°55'19" | 2617 | 14.5 | 0.28 | 3.67 | 112.85 | 5.94 | 177.20 | 17.80 | 0.71 | 332.18 | 24.6 | 0.2 | 146.0 | 7.88 |
| DT2&DT3<sup>b</sup> | 37°31'3" | 93°53'19" | 2617 | 14.5 | 103.51 | 6.57 | 0.23 | 13.18 | 180.97 | 22.14 | 0.27 | 358.30 | 137.0 | 0.1 | 8.30 | 7.94 |
| XT1&XT3<sup>b</sup> | 37°47'23" | 93°21'5" | 2576 | 13.0 | 10.73 | 0.30 | 0.14 | 0.60 | 17.87 | 1.75 | 0.17 | 36.83 | 37.3 | 0.5 | 4.60 | 7.02 |
| XT2 | 37°44'27" | 93°22'5" | 2576 | 15.0 | 0.11 | 0.09 | 2.74 | 0.30 | 0.44 | 0.80 | 0.16 | 20.09 | 0.7 | 0.1 | 52.00 | 7.00 |
| XT4 | 37°41'35" | 93°28'5" | 2576 | 15.0 | 0.12 | 0.10 | 3.17 | 0.36 | 0.51 | 0.83 | 0.18 | 28.46 | 0.5 | 0.2 | 47.00 | 7.01 |

Table1 Sampling record and Environmental conditions <sup>ac</sup>
| Sample name | N | E | H | T | Na<sup>+</sup> | K<sup>+</sup> | Ca<sup>2+</sup> | Mg<sup>2+</sup> | Cl<sup>-</sup> | SO<sub>4</sub><sup>2-</sup> | HCO<sub>3</sub><sup>-</sup> | TS | TN | TP | TOC | pH |
| --- | --- | --- | --- | --- | --- | --- | --- | --- | --- | --- | --- | --- | --- | --- | --- | --- |
| DT1 | 37°28'3" | 93°55'19" | 2617 | 14.5 | 0.28 | 3.67 | 112.85 | 5.94 | 177.20 | 17.80 | 0.71 | 332.18 | 24.6 | 0.2 | 146.0 | 7.88 |
| DT2&DT3<sup>b</sup> | 37°31'3" | 93°53'19" | 2617 | 14.5 | 103.51 | 6.57 | 0.23 | 13.18 | 180.97 | 22.14 | 0.27 | 358.30 | 137.0 | 0.1 | 8.30 | 7.94 |
| XT1&XT3<sup>b</sup> | 37°47'23" | 93°21'5" | 2576 | 13.0 | 10.73 | 0.30 | 0.14 | 0.60 | 17.87 | 1.75 | 0.17 | 36.83 | 37.3 | 0.5 | 4.60 | 7.02 |
| XT2 | 37°44'27" | 93°22'5" | 2576 | 15.0 | 0.11 | 0.09 | 2.74 | 0.30 | 0.44 | 0.80 | 0.16 | 20.09 | 0.7 | 0.1 | 52.00 | 7.00 |
| XT4 | 37°41'35" | 93°28'5" | 2576 | 15.0 | 0.12 | 0.10 | 3.17 | 0.36 | 0.51 | 0.83 | 0.18 | 28.46 | 0.5 | 0.2 | 47.00 | 7.01 |

### DNA sequencing

In this study, high-quality reads from DNA sequencing of the two lake samples were analyzed after strict quality control (Table S1). The results showed that the number of raw reads obtained for each sample in DT ranged from 85,776,056 to 88,722,652, while the number of clean reads after quality control ranged from 83,483,230 to 87,440,932. Compared with DT, in XT, the range of raw reads (84,456,600 to 105,040,790) and clean reads (82,990,342 to 103,270,840) was greater, and there were more reads overall. All samples were trimmed to obtain high-quality reads greater than 97% of the raw reads. After splicing, assembling, and predicting the sequences, a total of 1,677,194 ORFs were generated from the DT samples. After statistical analysis and prediction of the metagenomic sequencing data of four samples of XT, a total of 5,353,555 ORFs were obtained. The ORF data from the two salt lakes constitute a huge treasure trove of microbial genetic resources.

### Overview of microbial communities

Taxonomic classification of metagenomic data of the DT showed that 96.16% of the sequences corresponded to *Archaea*, while the other domains, *Bacteria* (2.71%), *Viruses* (1.06%), and *Eukaryota* (0.02%), accounted for less than 5% (Fig. S1A). In contrast, 93.09% of the sequences in XT corresponded to *Bacteria*, and the abundance of all other domains was less than 1% except for *Viruses* (4.38%). (Note: Relative abundances less than 0.01% in the next analysis are classified as “other.”)

### Taxonomic composition of bacterial communities

A summary of the bacterial distribution in the two salt lakes revealed that 33 phyla were present in both DT and XT (excluding candidate and unclassified *Bacteria*) (Table S2). The top five phyla in DT were *Proteobacteria* (61.84%), *Firmicutes* (8.81%), *Bacteroidetes* (8.48%), *Actinobacteria* (7.89%), and *Cyanobacteria* (2.29%) (Fig. 2A). *Proteobacteria* (65.59%) was the most abundant phylum in XT, followed by *Actinobacteria* (7.81%), *Bacteroidetes* (7.57%), *Verrucomicrobia* (4.14%), and *Planctomycetes* (4.05%).

**Figure 2.**
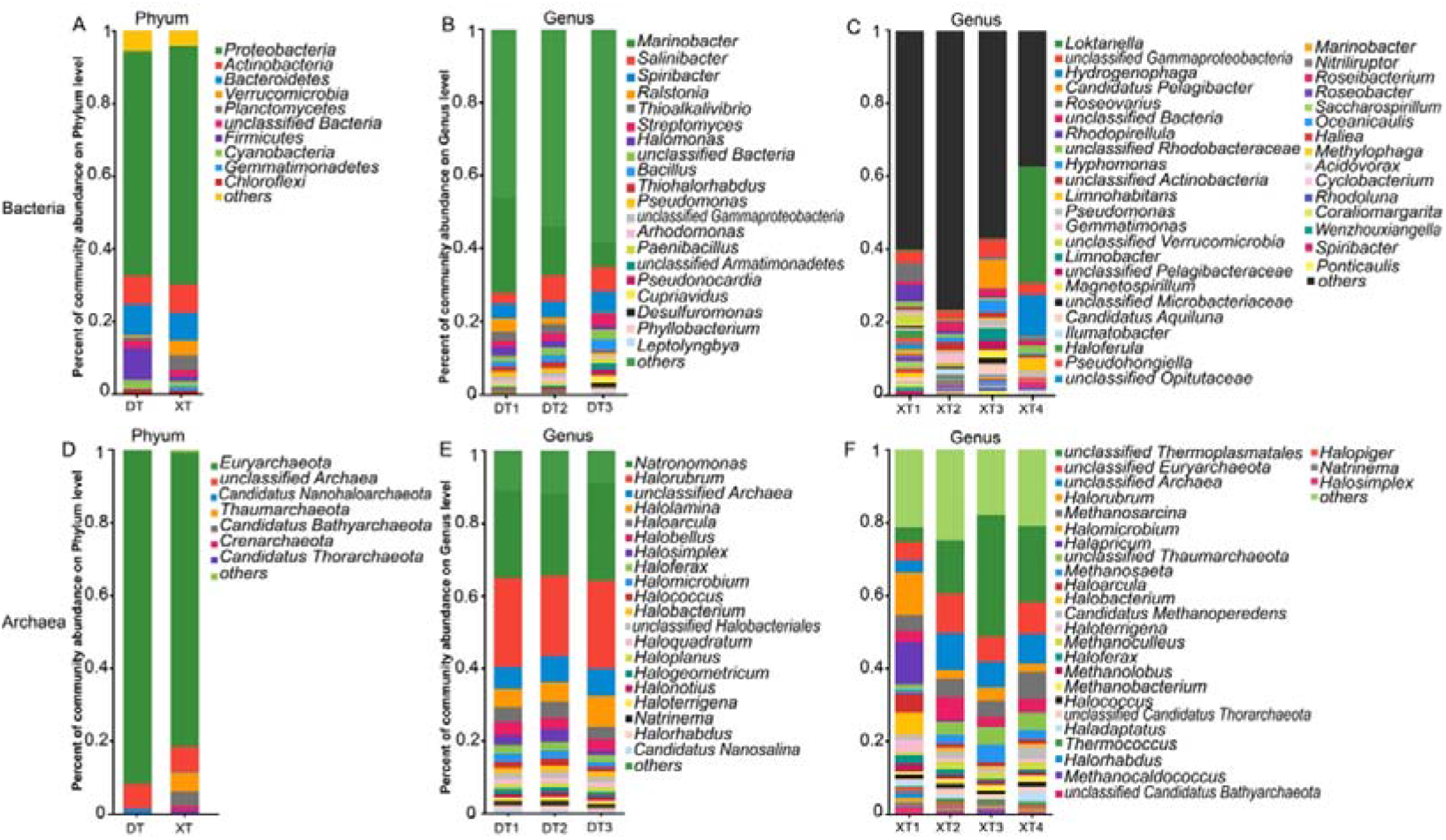
Taxonomic analysis of the DT and XT samples at the phylum and genus levels of *Bacteria* and *Archaea*.

With respect to bacterial genera, there were 1717 genera distributed in DT and 1906 genera in XT (Fig. S1B). Among these genera, *Marinobacter* (18.01%) was the most abundant genus in DT, followed by *Salinibacter* (5.09%), *Spiribacter* (4.43%), *Ralstonia* (2.35%), *Thioalkalivibrio* (2.18%), *Streptomyces* (2.06%), *Halomonas* (1.99%), and *Bacillus* (1.84%) (Fig. 2B). These dominant genera are mainly from the *Proteobacteria* phylum. Furthermore, the genera with relatively high abundances in XT were *Loktanella* (8.64%), *Hydrogenophaga* (3.17%), *Candidatus Pelagibacter* (2.03%), *Roseovarius* (1.71%), and *Rhodopirellila* (1.44%) (Fig. 2C). In this study, individual samples from the same salt lake converged consistently. For example, XT1–XT4 shared a similar genus population, and the dominant genus in each sample was *Roseovarius* (4.00%), *Gemmatimonas* (2.01%), *Candidatus Pelagibacter* (5.95%), and *Loktanella* (26.58%) respectively. However, except for *Marinobacter*, the abundance of the other genera in individual samples from DT was generally below 5%. At the species level, DT contained 8236 strains, XT had 12,621 strains, and 8203 strains were found in both lakes. The dominant species in DT and XT were *Salinibacter ruber* (5.09%) and *Loktanella vestfoldensis* (8.25%), respectively (Table S5).

### Taxonomic composition of archaeal communities

Pooled analysis of *Archaea* revealed four phyla were included in both DT and XT samples (excluding candidate and unclassified *Archaea*) (Fig. S1B). *Euryarchaeota* (91.48%) was the most abundant phylum in DT, and other phyla, including *Candidatus Nanohaloarchaeota* and *Thaumarchaeota*, were less than 5% (Fig. 2D; Table S6). The dominant phylum in XT was *Euryarchaeota* (80.63%), followed by *Thaumarchaeota* (5.18%) and *Candidatus Bathyarchaeota* (3.40%).

At the genus level, there were also great differences between the two lakes, and 153 archaeal genera in DT and 149 genera in XT were identified. The major genera in DT were *Halorubrum* (23.69%) and *Natronomonas* (24.61%) (Fig. 2E). As a member of *Natronomonas, Natronomonas moolapensis* (22.33%) was the dominant strain of DT (Table S9). Compared with XT, *Halomicroarcula*, *Haloarchaeobius*, *Halomarina*, *Haloparvum Halohasta*, *Salinigranum*, *Halosiccatus*, and *Natronoarchaeum* were unique genera in DT. The top four genera in XT were *Thermoplasmatales* (17.01%), *Euryarchaeota* (7.62%), *Halorubrum* (5.74%), and *Methanosarcina* (4.81%) (Fig. 2F). Other genera, including *Nanoarchaeum*, *Acidianus*, *Candidatus Thalassoarchaea*, and *Methanosarcinales*, were distinct in XT. The *Thermoplasmatales* archaeon (14.05%), a member of the *Thermoplasmatales* genus, was the dominant strain in XT. In addition, *Methanosarcina barkeri* was predominantly present in XT (Table S9), which was previously studied in the metabolic regulation of ammonium inhibition (27).

### Functional annotation of clusters of orthologous groups (COG)

A visual analysis of COGs at the category level shows the abundant genes involved in the fundamental metabolism of the DT and XT communities (Table S10). At the functional level, a total of 17 different COG categories were identified (gene abundance <0.01 was classified as other) (Fig. 3A). Among them, amino acid transport and metabolism (E), replication, recombination, and repair (L), energy production and conversion (C), and inorganic ion transport and metabolism (P) were present at a relatively high proportion (accounting for >5%) (Table S11). It is noteworthy that the cell wall/membrane/envelope biogenesis (M) function abundance in XT was higher than that in DT. The highest abundance was functional unknown (S), suggesting that the potential functional resources of these two salt lakes deserve further exploration.

**Figure 3.**
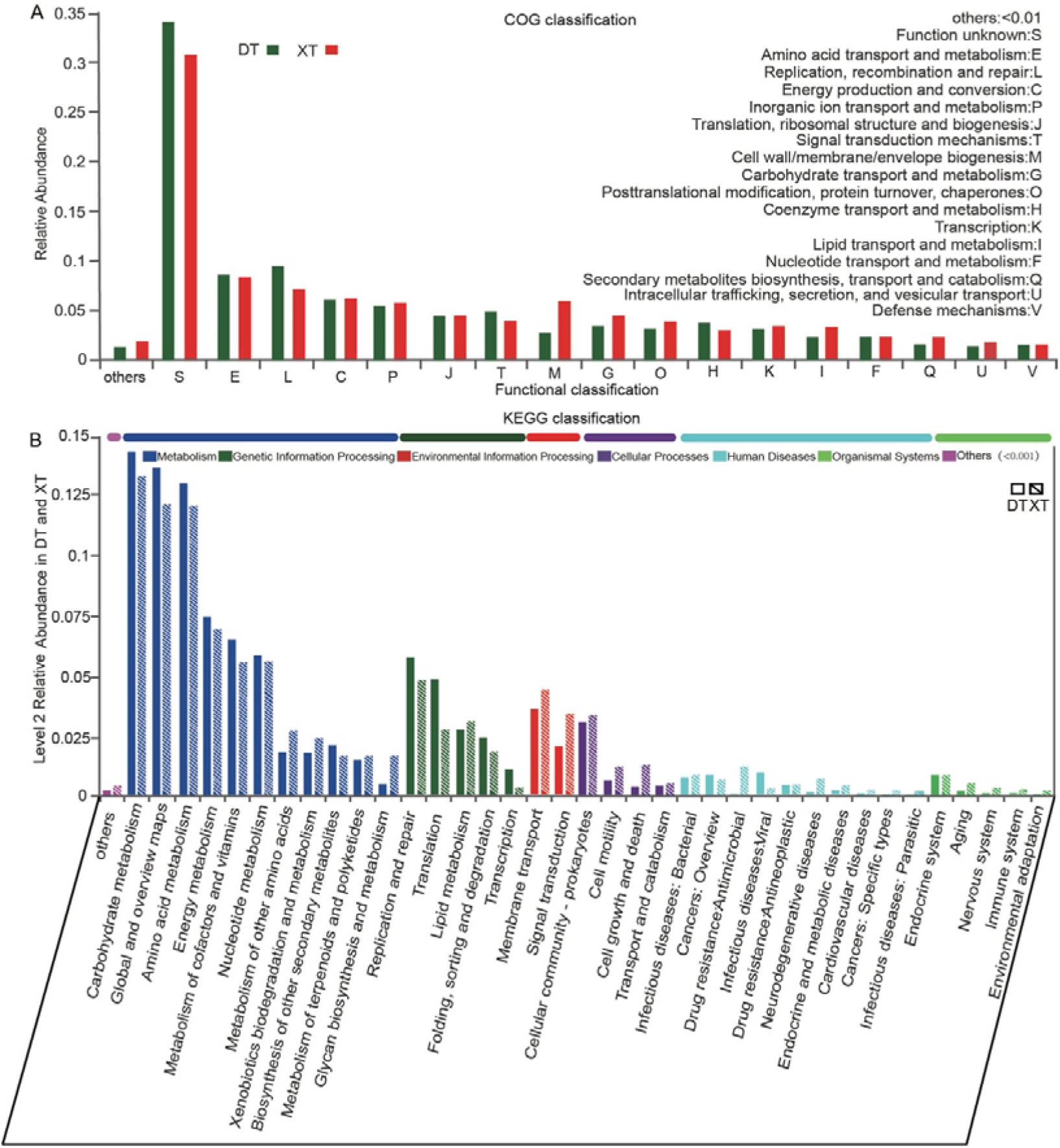
COG and KEGG function annotation of the DT and XT communities. (A) The x-axis represents the functions in the COG classification, the y-axis represents the proportion of each function in the sample, and different colors represent the two lakes. (B) The x-axis indicates the KEGG metabolic pathways at classification level 2, the y-axis indicates the corresponding percentages of the samples, and different colors show the clustering results.

### Functional annotation of the KEGG metabolic network

The KEGG metabolic network showed that the high abundance genes were involved in metabolism, followed by genetic information processing (Fig. 3B). The significantly annotated metabolism categories in DT included carbohydrate metabolism, global and overview maps, amino acid metabolism, energy metabolism, metabolism of cofactors and vitamins, and nucleotide metabolism. However, the metabolism categories of other amino acids, xenobiotics biodegradation and metabolism, metabolism of terpenoids and polyketides, and glycan biosynthesis and metabolism were slightly more annotated in XT than in DT. Some gene information processing categories, including replication and repair; translation; and folding, sorting, and degradation, were more annotated in DT, while lipid metabolism was relatively more annotated in XT. Furthermore, cellular processes and environmental information processing, including membrane transport, were more annotated in XT. In this study, human diseases and organismal systems were rarely noted. Multiple mechanisms may be involved in adaptation to extreme environments but to different degrees.

### Major metabolic pathways and halophilic mechanisms

The mechanisms of microbial halophiles mainly included 1) the regulation of Na^+^ efflux- and K+ uptake-related genes (part of the so-called “salt-in” strategy), and 2) organic compatible solute biosynthesis and transport (accumulation of compatible solutes by uptake or de novo synthesis) (28, 29). Han et al. demonstrated through transcriptome analysis that the glycine, serine, and threonine metabolic pathways (ko00260) and the ABC transporter (ko02010) play important roles in salt-stressed environments (30). We used KEGG gene annotations to study compatible solute transport and the biosynthesis pathways of microorganisms in a hypersaline salt lake (DT) and moderate salt lake (XT).

In this study, ko00260 was the predominant metabolic pathway in the analysis of amino acid metabolic pathways, followed by alanine, aspartate, and glutamate metabolism (ko00250), and cysteine and methionine metabolism (ko00270) (Fig. S2A). Eight metabolic pathways that were significantly different are listed in Fig. 5A. The results showed that the biosynthetic pathways ko00400 (phenylalanine, tyrosine, and tryptophan biosynthesis), ko00220 (arginine biosynthesis), ko00300 (lysine biosynthesis), and ko00290 (valine, leucine, and isoleucine biosynthesis) were significantly upregulated in DT. In contrast, the pathways ko00380 (tryptophan metabolism), ko00360 (phenylalanine metabolism), ko00340 (histidine metabolism), and ko00310 (lysine degradation), which are related to metabolism and degradation, were significantly upregulated in XT. Similarly, the ko00260 metabolic network of enzymes showed differences between DT and XT (Fig. S3). D-amino-acid oxidase (EC: 1.4.3.3), guanidinoacetate N-methyltransferase (EC: 2.1. 1.2), and serine racemase (EC: 5.1.1.18) were the main enzymes significantly enriched in XT, while alcohol dehydrogenase (EC: 1.1.1.1) was significantly enriched in DT (Fig. S4).

Most bacteria that synthesize ectoine and hydroxyectoine are isolated from extreme environments, such as low temperature, dryness, or high salinity (31). Discrepancies in salinity might lead to differences in solute synthase expression. Therefore, the enzymes involved in ectoine/hydroxyectoine synthesis were also evaluated at the KEGG orthology level in this study (Fig. 4B, Fig. S2B). Increased salinity may reinforce the progress of the biosynthesis of amino acids, carbon metabolism, glycolysis/gluconeogenesis, DNA replication, and the TCA cycle pathways in response to extreme stress (Fig. S2C). Therefore, compatible solutes and salt-in strategies were the preferred contingency strategy of bacteria in DT and XT.

**Figure 4.**
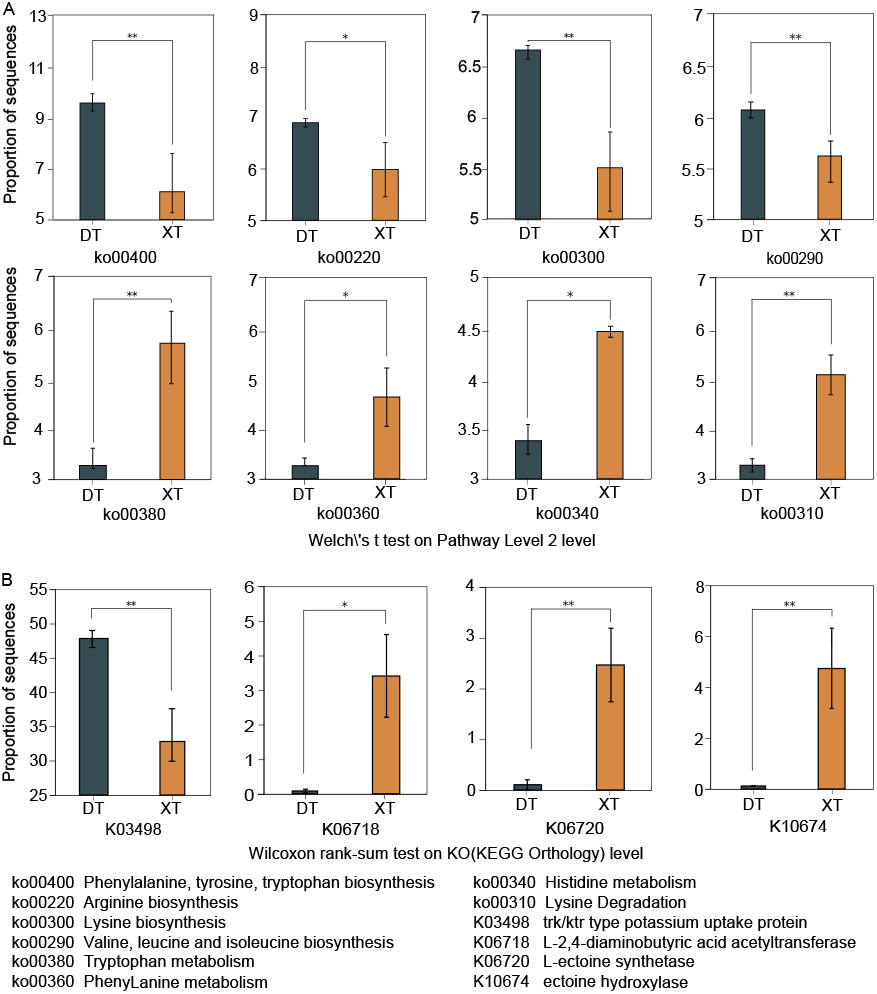
(A) Eight pathways with the most significant differences at level 2 of KEGG; (B) KEGG orthology with significant differences in the ko00260 pathway; 0.01 < p ≤ 0.05* and 0.001 < p ≤ 0.01 **.

There was a significant abundance of K03498-related *trkH*, *trkG*, *ktrB*, and *ktrD* genes in the DT samples for potassium ion uptake (32). Therefore, when the concentration of exogenous salt increases, microorganisms in DT may respond to salt stress by upregulating the expression of trk/ktr potassium uptake proteins (K03498) (Fig. 4B). Notably, ectoine hydroxylase (K10674) and L-ectoine synthetase (K06720) were upregulated in XT. This promoted the synthesis of the compatible solute ectoine and hydroxyectoine in response to extreme environmental stresses. Different mechanisms may have positive effects on the survival of microorganisms in DT and the active metabolism and transformation of microorganisms in XT.

### Association analysis of dominant genera and environmental factors

Hypersalinity in combination with intense UV irradiance, high lithium concentrations, and moderately acidic pH makes unique brines for studies on diversity and the adaptation of polyextremophilic microorganisms (33). To illustrate the associations between community composition and the environmental parameters, detrended correspondence analysis (DCA) was first used to identify axis lengths > 3, after which CCA was used to detect the key factor interactions of the dominant genera (Table S14). The correlation R^2^ and p-values of the analysis results are listed in Table S15.

The most abundant (relative abundance >5%) bacterial genera across the seven samples were *Marinobacter, Salinibacter*, and *Spiribacter*. In addition, the most abundant archaeal genus was *Natronomonas*, and other *Archaea* were found in individual samples, including *Halorubrum* and *Methanosarcina*. The results of the CCA showed that the total phosphorus (TP) content was most positively correlated with *Marinobacter*, followed by *Spiribacter*, *Roseovarius*, *Hyphomonas*, *llumatobacter*, and *Gemmatimonas* (Fig. 5). Furthermore, *Salinibacter* correlated with TOC content. It is worth noting that other dominant genera, including *Natronomonas* and *Halorubrum*, and some archaeal genera from *Euryarchaeota* phylum were strongly correlated with pH, TN, Na^+^, Cl^−^, HCO_3_^−^, and Ca^2+^. In addition, these representative genera were highly positively correlated with TS, K^+^, and Mg^2+^ levels. In general, the interaction of dominant genera and the environment was completely different in the two salt lakes. The dominant genus in DT was positively correlated with the total salt, which mainly consisted of sodium, chlorine, magnesium, and sulfate ion concentrations. Although XT4 was located far from the other three samples, the samples in XT were mainly positively correlated with *Bacteria*, and TP was the determinant factor.

**Figure 5.**
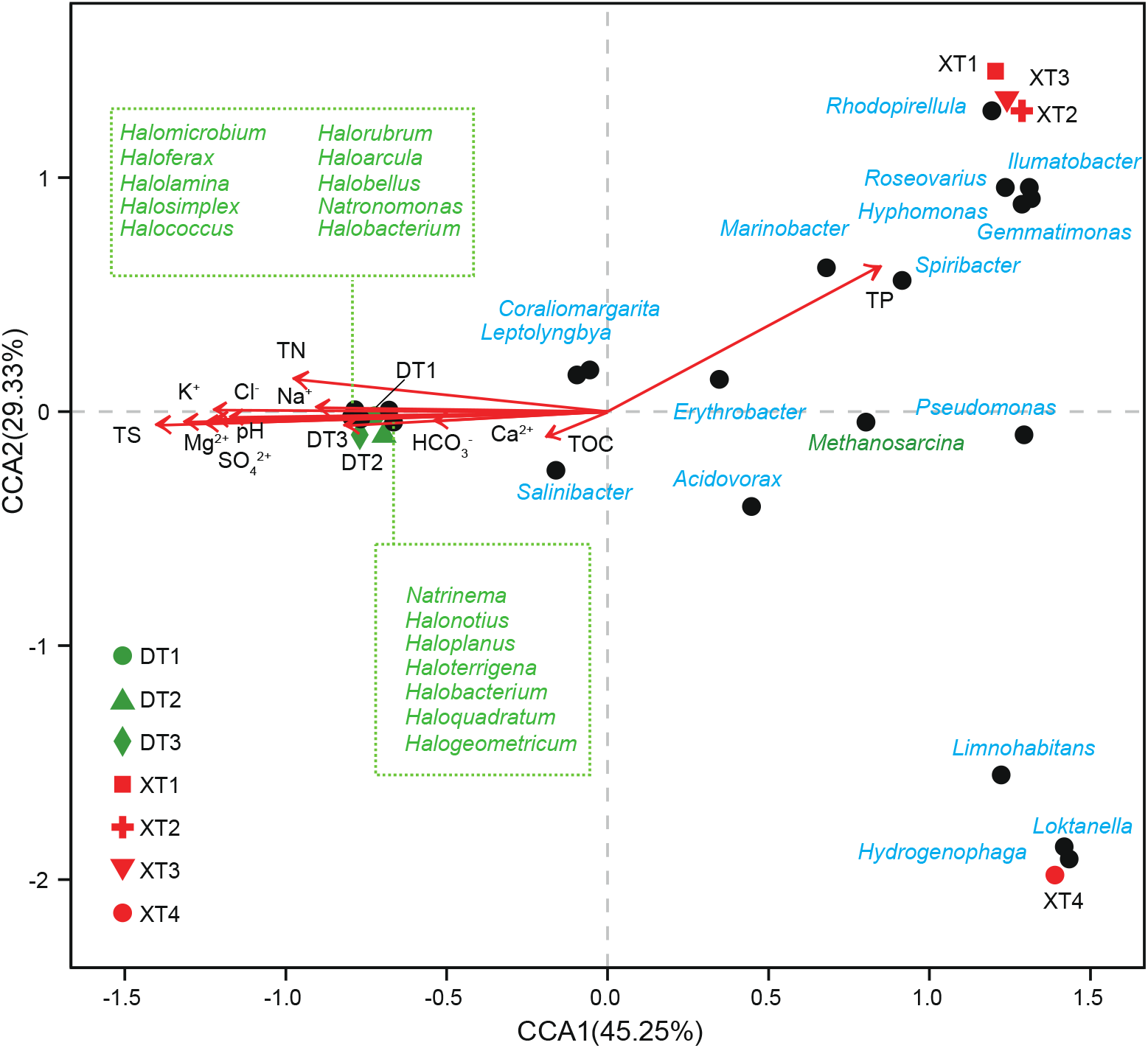
CCA of the relationship among dominant genera and physicochemical variables in the two salt lakes. The sample coordinate table shows the relative positions of samples in each dimension after dimension reduction. Environmental factors are represented by arrows in red, and the length of the arrow line represents the magnitude of the correlation. The longer the line, the greater the correlation, and vice versa. The black dots in the figure represent the genus of *Bacteria* and *Archaea* with names in different colors. Green is *Archaea* and blue is *Bacteria*. Different colors and shapes represent different subgroups.

## DISCUSSION

### Analysis of environmental characteristics

Salinity is an important factor in determining the type of a lake. According to Zheng et al. (4), the salinity of DT and XT were 332.0 and 338.0 g/L, respectively in 2010; therefore, they were both classified as hypersaline lakes (34). Although the salinity of DT did not alter notably in 2020 (34) when compared with previous years, the salinity of XT dropped to 124.8 g/L in the same year (35). In this investigation, DT (with a salinity of 332–358 g/L) remained regarded as a hypersaline lake (12), while XT became regarded as a moderate saline lake with a salinity of 20–36 g/L (Table 1) (36).

Evaporation and seepage are two major factors altering the salinity of lakes. In addition, river water, groundwater and brine play roles in maintaining stable water levels and salinity (7). However, their interactions vary in wet and dry seasons. The size of the Taijinar Salt Lake was reduced significantly in 2015 owing to evaporation, and the original water-salt balance changed after 2020 (37, 38). The NLGL supplies fresh water to the XT and results in the salinity decreasing. The main ions fluctuate as a result of these combined effects (Table 1). The pH of DT and XT in this investigation was weakly basic. As a result, we hypothesized that global warming may also be responsible for the decline in the salinity of lakes on the Tibetan Plateau (39).

### Microbial community structure in DT and XT

In this study, the microbial composition of XT was dominated by *Bacteria* (93.09%), while the most abundant microbe in DT was *Archaea* (96.16%) (Fig. 2A). The change in microbial species compositions in salt lakes with salinity is characteristic (14, 40). *Archaea* prefers high-salt and high-temperature environments(41). The archaeal communities in hypersaline waters are common, which was confirmed by Lina et al. (42). For example, the sediment communities of the deep lakes of the Antarctic are almost entirely composed of halophilic *Archaea* (43). Even in Salar de Uyuni, Bolivia, only the archaeal *Halobacteria* taxa were founded, while bacterial taxa were completely absent (33). Although there are some habitats with their own unique and infrequent microbial community distributions (44, 45), microbial community structure and diversity in lake ecosystems are mainly affected by salinity (ion composition), followed by pH, nutrition, and temperature (34, 46). In this study, salinity was the primary significant factor driving differences in species composition.

### Bacterial phyla and genera in DT

According to the results, *Proteobacteria* (61.84%) was the most prevalent phylum in DT, followed by *Firmicutes* (8.81%), *Bacteroidetes* (8.48%), *Actinobacteria* (7.89%), and *Cyanobacteria* (2.29%) (Fig. 2A). Although bacterial sequences accounted for only 2.72% of all sequences, there were still several dominate genera in DT (Fig. 2B; Table S4). For example, *Marinobacter* (18.01%) in DT was the most prevalent genus and positively correlated with TP content (Fig. 5). Previous studies demonstrated its ability to synthesize various compounds and degrade hydrocarbons in high-salinity environments (47). In addition, *Marinobacter* were shown to control the formation of biofilms associated with calcium levels(48). *Halomonas*, *Salinibacter*, and *Salicola* genera were at a higher abundance in DT, of which *Salicola* was unique. *Salicola* resists UV-B irradiation at the optimum salinity (NaCl) for its growth, and its member *Salicola marasensis* can accumulate glycine betaine at a wide range of salinity (10–25%) (49, 50). Furthermore, previous studies showed that the average lithium content in the dry salt flats of DT was 0.8 g/L (115.26 mmol/L) (5). High lithium concentrations (50–680 mmol/L) may have inhibitory effects on extreme halophilic bacteria including *Halomonas*, *Salinibacter*, and *Salicola* (33, 51). However, the interaction between the rich lithium concentration and extreme halophilic bacteria in DT needs to be further explored.

### A unique and diverse bacterial structure in XT

The bacterial community analysis of XT found that *Proteobacteria* (65.59%) was the dominant phylum, followed by *Actinobacteria* (7.81%) and *Bacteroidetes* (7.57%) (Fig. 2A). *Loktanella* (8.64%), as the dominant genus of XT, may solve the problem of co-utilization of complex carbon and nitrogen sources (Fig. 2C) (52). The presence of *Poribacteria* is consistent with the report that most moderate and extreme halophilic bacteria exist in the subgroups of *Proteobacteria* and *Firmicutes* (53).

Some low-abundance genera may also play a prominent role in the ecology of salt lakes. Based on genome sequence reconstruction, unique functional features of *Candidatus Poribacteria* were identified by Sheila et al. (54). Among them, we were most interested in its metabolism of the osmolytes, including ectoine and glycine betaine, and cellulosome anchoring of extracellular enzyme complexes through cohesin and dockerin domains. Some samples also contained unique genera that perhaps work together with dominant genera in the region to maintain ecosystem stability. *Hydrogenophaga* is a genus that was mainly distributed in XT4, and its members play important roles in many fields. For example, *Hydrogenophaga sp.* strain PBC is an effective degrader of 4-aminobenzenesulfonate, which may have strong potential in biological metabolism and degradation (55). Sebastian et al. investigated *Hydrogenophaga pseudoflava* as a novel host for producing chemicals from syngas (56). They also indicated that it could be a promising candidate for future aerobic production processes. Furthermore, *Haloplasma* from the deep-sea anoxic brine lakes is a member of *Haloplasmatales* and was relatively rich in the XT4 sample. The presence of these genera suggests that the salt lake microbial community may have a strong ability to maintain a stable ecosystem in XT. However, the linkages and transformations of their interaction networks during the evolutionary process need further in-depth study.

### Archaeal community and dominant genus in DT and XT

Microbial communities in lakes on the Tibetan plateau did not change with geographical factors but strongly correlated with the physicochemical parameters of each lake (57). The salinity was the major driver for outlining microbial community structure. In this study, the dominant archaeal phylum in both DT and XT was *Euryarchaeota*. It is known that higher salinity may have facilitated *Euryarchaeota* survival and growth (58), although this may result in low diversity (59). Microorganisms living in hypersaline lakes must endure salinity stress (60), and therefore, only a few microorganisms can survive in such an extremely harsh environment. This may be the reason for *Archaea* having a lower diversity but greater abundance in DT. This was demonstrated by the fact that *Archaea* accounted for 91.48% of the DT microbial community (Figure 2A), but the number of phyla, orders, and genera were not very high (Table S6–7). Moreover, the high quality of DNA sequences finally obtained in DT and XT illustrate this point (Table S1).

Halophilic archaea counteract this salinity stress mainly through a salt-in strategy (57, 61). Among the *Archaea* studied so far, organisms using this strategy are divided into two groups: the majority of organisms maintain osmotic balance by accumulating K+ in the cell and releasing Na+ outside of the cell, while a few organisms use inorganic ions almost exclusively to balance a relatively constant internal K+ (41). For these few *Archaea*, the counteraction of salinity and the intracellular environment, as well as the interaction of nonionic osmolytes with compatible solutes, together decide the mechanism employed against osmotic stress, with salinity being the key factor. Furthermore, members of Euryarchaeota are involved in metabolism related to sulfur, carbon, and nitrogen to maintain stable ecosystem functions in high-salinity lakes such as DT and XT (62–64). *Candidatus Nanohaloarchaeota* and *Thaumarchaeota* were also among the top five dominant *Archaea* in this study. Among them, *Thaumarchaeota* was also identified during the microbial diversity analysis of Gahai Lake and Xiaochaidan Lake (58). *Nanohaloarchaeota* members are dominant in high-salinity environments (especially in a salinity exceeding 200 g/L).

We noted a high abundance of *Halorubrum* and *Natronomonas* in our samples. There is a symbiotic relationship between *Halorubrum lacusprofundi* and *Candidatus Nanohaloarchaeum antarctic* (58). The strategies adopted by these unique members under extreme conditions may contribute to the survival of related microorganisms in DT.

### Microbial metabolic network patterns by salinity

The geochemical regime of different sites within extreme environments exerts a clear selective force on microbial communities and on patterns of microbial activity (65). The adaptation of microbial communities to face fluctuating salinity may be achieved by regulating the relative abundance and function of some taxa (13). However, bacterial growth and metabolism in DT may be restricted by high salinity in MgSO_4_ subtype lakes, and the diversity was also low. This hypothesis, low diversity under high-salinity conditions, was demonstrated by the fact that the number of ORFs obtained from the DT samples was generally lower than those of XT (Table S1). Similarly, microbial adaptation under a changed salinity may be achieved by adjusting certain community functions, such as regulation of membrane transport proteins, biosynthesis, and metabolic genes. GO and KEGG enrichment analyses indicated that amino acid transport and metabolism may play a major role in osmotic adaptation induced by moderate and high salinity (Fig. 4). Interestingly, it appears that microorganisms involved in amino acid metabolism and carbon metabolism were more active in DT (hypersaline lake) than in XT (moderate saline lake) (Fig. 4). However, regarding microorganisms in DT, high-salt stimulation inhibited membrane transport and lipid metabolism, which are important for maintaining microbial life survival. The regulation of membrane proteins may affect the survival of prokaryotes under hyperosmotic stress and lead to the specific structure of *Haloarchaea* (66, 67).

In this study, glycine, serine, and threonine metabolism (ko00260) accounted for a similar proportion of the total amino acid metabolic pathway in microorganisms from both salt lakes, 13.98% in DT and 13.22% in XT (Fig. S2A). However, microorganisms in DT and XT have their own dominant metabolic pathways for their respective environments. For example, metabolism-related pathways such as ko00400, ko00220, ko00300, and ko00290 were prominently annotated in DT. In contrast, ko00380, ko00360, ko00340, and ko00310 were significantly annotated in XT. This was consistent with its classification and results in terms of microbial composition and function.

There were different adaptation strategies in DT and XT against the extreme environment. First, genes associated with the synthesis of compatible solutes, including ectoine (K06720) and hydroxyectoine (K10674), were significantly annotated in XT (Fig. 4B). The synthesis of these compatible solutes and their related pathways provide a suitable opportunity for varied life forms in XT. However, microorganisms in DT tended toward enhanced basal metabolism and upregulated expression of trk/ktr-type potassium uptake proteins to generate a salt-in adaptation strategy in response to salt stress. Notably, this may have contributed to the low pathogenicity and limited secondary metabolite synthesis of microorganisms in DT. Second, low or moderate halophilic bacteria use intracellular synthesis or uptake of compatible solutes as their main coping mechanism (57, 61). In contrast, most halophilic *Archaea* (even nonhalophilic *Archaea*) respond to higher osmotic pressure by uptake of K^+^ and export of Na^+^ (32, 41). Therefore, this salt-in adaptation strategy allows halophilic *Archaea* to thrive and dominate in extreme hypersaline environments (57, 60). The basic mechanism behind this salt-favored strategy is the intracellular enzyme machinery of halophilic *Archaea*, such as *Halobacterium* and *Haloferax*, that requires a certain molar concentration of salt to maintain its proper conformation and activity(68). *Bacteria* and *Archaea* may exist in direct competition that results in halophilic *Archaea* taking the energetic advantage in carbohydrate metabolism, amino acid metabolism, energy metabolism, and metabolism of cofactors and vitamins, eventually dominating in high-salinity lakes (57). In addition, changes in physical and chemical properties explain differences in the microbial community composition and predict microbial functions in salt lakes. In summary, the environment mainly influences the overall function of DT and XT microorganisms and thus causes the differences and specificity of the community structure. The microbial structural diversity not only adapted to the specific environment but also formed a complex network of interactions with differential factors that together built the unique ecosystems of these two salt lakes.

Although we have used metagenomic sequencing technology to initially explore the mystery of salt lakes on the Tibetan Plateau, the changes in microbial abundance and species caused by environmental changes, such as seasonal variations in inputs, temperature, and precipitation, are still unknown. The preliminary analysis of the general situation of the two salt lakes in this study may provide a basic reference for the profound interactions of microbial differences and environments in the salt lakes on the Qinghai-Tibet Plateau under different conditions.

### Conclusions

The microbial composition of saline lakes is quite complex (especially in hypersaline lakes), and the diversity of *Archaea* in extreme environments and their ecological functions have boomed in recent years. To the best of our knowledge, this study is the first discussion on the microbial structural composition of DT and XT based on a metagenomic analysis. The two salt lakes are closely located with different salinity and other factors. However, the microbial communities of the two salt lakes are completely different. Each sampling site was dominated by a few specific genera. We believe that salinity was a key factor affecting species distribution and microbial function in the two salt lakes. Moreover, the interaction network of environmental characteristics and the microbial community composition may coordinate with a high-salinity environment and maintain saline ecosystem stability. Their adaption functions in a harsh environment were explored, and these results will expand our knowledge of the biodiversity of saline lakes on the Qinghai-Tibet Plateau.

## MATERIALS AND METHODS

### Sample collection and physicochemical analysis

A mixture of sediment and water was collected from DT and XT during the summer of 2020 (July 20) (Fig. 1). Two-liter turbid water samples were obtained 2 m from the lakeshore at a 15 cm depth from the water surface at each site. The temperature, pH, altitude, latitude, and longitude of the water were recorded during sampling. After removing impurities such as sand and stone, the pretreated solution was treated using a 0.45-μm polyether sulfone acetate membrane (Millipore, USA) to obtain filtrate. The residue was immediately transferred to a −80°C freezer for further DNA extraction. The filtrate was stored at 4°C and sent to Micor-Analysis Technology Co., Ltd. for physicochemical analysis.

### Metagenomic DNA extraction and sequencing

Total genomic DNA was extracted from all the collected samples using the E.Z.N.A.® Soil DNA kit (Omega Bio-tek, Norcross, GA, USA) according to the manufacturer’s instructions. The concentration and purity of the extracted DNA were determined with a TBS-380 and NanoDrop 2000, respectively. The extracted DNA quality was analyzed using a 1% agarose gel and then was fragmented using a Covaris M220 (Gene Co. Ltd., Shanghai, China) with an average size of approximately 400 bp. Paired-end sequencing libraries were prepared using NEXTFLEX™ Rapid DNA-Seq (Bioo Scientific, Austin, TX, USA). Paired-end sequencing was performed on an Illumina Novaseq 6000 system (Illumina Inc., San Diego, CA, USA) using NovaSeq reagent kits according to the manufacturer’s instructions (www.illumina.com).

The raw data sequences were trimmed to obtain high-quality short reads using Fastp (https://github.com/OpenGene/fastp, version 0.20.0) by removing low-quality and ambiguous reads (reads with unknown nucleotides “N”) (20). The high-quality short reads with contigs ≥ 300 bp were assembled using Megahit software (https://github.com/voutcn/megahit, version 1.1.2) (21). Subsequently, MetaGene (http://metagene.cb.k.u-tokyo.ac.jp/) was used to predict the open reading frames (ORFs) of the assembled contigs for functional annotation. Non-redundant gene catalogs were clustered and constructed using CD-HIT (http://www.bioinformatics.org/cd-hit/, version 4.6.1) with 90% sequence identity and 90% coverage (22). High-quality reads from each sample were aligned to the non-redundant gene catalogs separately using SOAPaligner software (http://soap.genomics.org.cn/, version 2.21) to calculate the gene abundance with 95% identity (23).

### Bioinformatic analysis

An E-value ≤ 1e^−5^ was used during the blastp operation. Taxonomic and functional annotation information was obtained by comparing representative sequences of non-redundant gene catalogs with NR databases, the Evolutionary Genealogy of Genes: Non-supervised Orthologous Groups database (EggNOG, http://eggnog.embl.de/) (24), and the Kyoto Encyclopedia of Genes and Genomes database (KEGG, http://www.genome.jp/kegg/) (25) using DIAMOND (http://www.diamondsearch.org/index.php, version 0.8.35) (26).

### Statistical analysis

The species annotation method was best-hit, and Bray-Curtis dissimilarity was used as a distance algorithm. The differences between different groups were calculated using Welch’s test (Welch’s inverted confidence 0.95). Differential enzyme testing and visualization based on KEGG annotation results used a two-sided Wilcoxon test (*p* < 0.05 for significance) for specific pathways. Redundancy analysis (RDA)/canonical correspondence analysis (CCA) was used to obtain regression R^2^ values and p-values to further investigate the relationship between environmental characteristics and the sample.

### Data availability

The raw Illumina sequencing data was uploaded to the NCBI database archive with the accession number PRJNA984434 (https://www.ncbi.nlm.nih.gov/sra/PRJNA984434).

## Acknowledgments

This work was supported by the National Natural Science Foundation Project of China (21967018) and National Natural Science Foundation Project of China (32260019).

## Authors’ contributions

Man Zhang analyzed the data and wrote the manuscript. Yongzhen Li ensured the accuracy or integrity of the work. Yongzhen Li and Derui Zhu designed the experiments and supervised the revision of the manuscript. Qifu Long and Guoping Shen collected the experimental samples. Jiangwa Xing analysed the data. All authors edited the manuscript and approved the final draft.

